# Predictability shifts from local to global rules during bacterial adaptation

**DOI:** 10.1101/2022.05.17.492360

**Authors:** Alejandro Couce, Melanie Magnan, Richard E. Lenski, Olivier Tenaillon

## Abstract

The distribution of fitness effects of new mutations is central to predicting adaptive evolution, but observing how it changes as organisms adapt is challenging. Here we use saturated, genome-wide insertion libraries to quantify how the fitness effects of new mutations changed in two *E. coli* populations that adapted to a constant environment for 15,000 generations. The proportions of neutral and deleterious mutations remained constant, despite large fitness gains. In contrast, the beneficial fraction declined rapidly, approximating an exponential distribution, with strong epistasis profoundly changing the genetic identity of adaptive mutations. Despite this volatility, many important targets of selection were predictable from the ancestral distribution. This predictability occurs because genetic target size contributed to the fixation of beneficial mutations as much as or more than their effect sizes. Overall, our results demonstrate that short-term adaptation can be idiosyncratic but empirically predictable, and that long-term dynamics can be described by simple statistical principles.

**One-Sentence Summary:** Couce et al. demonstrate that short-term bacterial adaptation is predictable at the scale of individual genes, while long-term adaptation is predictable at a global scale.

## Main Text

Evolution in asexual populations results from the accumulation of new mutations. Therefore, detailed knowledge of the proportions of mutations that are beneficial, neutral, or deleterious is important for predicting the course and outcomes of evolution. Indeed, assumptions about the distribution of fitness effects (DFE) of new mutations lie at the core of many theories describing fundamental evolutionary phenomena, including the speed of adaptation (*1*), fitness decay in small populations (*2*), the maintenance of genetic variation (*3*), the probability of parallel (*4*) versus divergent (*5*) evolution, the pace of the molecular clock (*6*), and the evolution of sex (*7*) and mutation rates (*8*). Driven by this interest, many experimental and comparative studies have produced estimates of this distribution in different organisms. While most studies have been small in scale (*9*) or focused on narrow genomic regions (*10*), some important features appear similar across a variety of model systems. In particular, most mutations are neutral or deleterious, lethal mutations form a distinct class within the deleterious tail, and those rare beneficial mutations are usually exponentially distributed (*11*).

However, the DFE only indicates what is possible at a particular point in time, and it is unknown how long the distribution will remain relevant as evolution proceeds and especially as beneficial substitutions accumulate. Predictions on how the shape of the DFE changes are implicit in some theoretical models, most famously that the beneficial tail should approach an exponential distribution near a fitness peak (*12, 13*). The picture is more ambiguous for the deleterious tail, with different models predicting it to become heavier (*14*) or lighter (*15*) with adaptation. In any case, these predictions address only the macroscopic form of the DFE, with little attention to the microscopic processes underlying the changes in shape. And while the shape influences the dynamics, the microscopic details—especially the sign and intensity of interactions among mutations (*i*.*e*., epistasis)—determine the outcomes of adaptation (*16*). For example, in the absence of epistasis, adaptation will shorten the beneficial tail simply by the process of sampling without replacement. In this case, a complete DFE would suffice to specify the probabilities of all possible adaptive trajectories in a given environment. At the other extreme, if sign epistasis is the norm—such that mutations go from beneficial to deleterious, and vice versa, as the genome evolves (*17*) —then new mutations will continually change the effects and rank order of the remaining mutations, rendering futile any prediction about adaptive trajectories beyond the very short term.

### High-throughput insertion mutagenesis and fitness measurements

To determine experimentally how adaptation and epistatic interactions change the DFE, one would ideally like to measure the relative fitness of a large set of mutants at multiple time points along a broad and well-characterized adaptive trajectory. To do so, we take advantage of the long-term evolution experiment (LTEE) in which populations of *Escherichia coli* have been adapting to a glucose-limited medium for tens of thousands of generations, resulting in large fitness increases (*18*). Examples of both weak and strong epistasis among beneficial mutations have been reported in this system (*16, 19*). Moreover, the most important mutations driving adaptation have been identified from signatures of parallelism in whole-genome sequences (*20, 21*), allowing the predictive capacity of a DFE at one time point to be compared with the actual fate of mutations during later adaptation. To measure the DFE, we created genome-wide libraries of insertion mutants using a transposon engineered to capture the 14-bp sequence adjacent to each insertion site, which in most cases identifies unequivocally the target locus (*22*) (Methods, Fig. S1). We typically identified >100,000 different insertion mutants, which mapped to >78% of the ancestral genome’s 8424 loci including both open reading frames (ORFs) and intergenic regions (Methods).

We estimated the fitness effects of all these mutants as selection coefficients, obtained by tracking the frequency trajectory of every allele during 5-day, bulk competition assays under the same conditions as in the LTEE (Fig. 1, a-c; Methods). Although transposon insertions typically cause losses of function, we also saw two types of more subtle effects (Fig. 1, d). First, an insertion in the C-terminus of a gene may cause only a partial loss of function or even a change in function. This outcome is prominently revealed by the tolerance of many essential genes to insertions in that region, notably including insertions in *topA* and *serB* that show large benefits (*23*). Second, the position of many beneficial insertions, including in intergenic regions and genes upstream of known targets of adaptation in the LTEE, suggests changes in gene expression. Polar effects within transcription units are expected because the 1.5 Kb insert carries two transcriptional terminators after a kanamycin resistance gene. To detect these phenomena, while also ensuring robust fitness estimates, we divided each locus into 5 segments of equal length and then pooled all insertions in each segment. As an added benefit, comparing the fitness effects among segments of the same locus allows identification of potential artifacts and provides an internal control to quantify the reproducibility of the fitness estimates (Methods, Fig. S2).

**Fig. 1.**
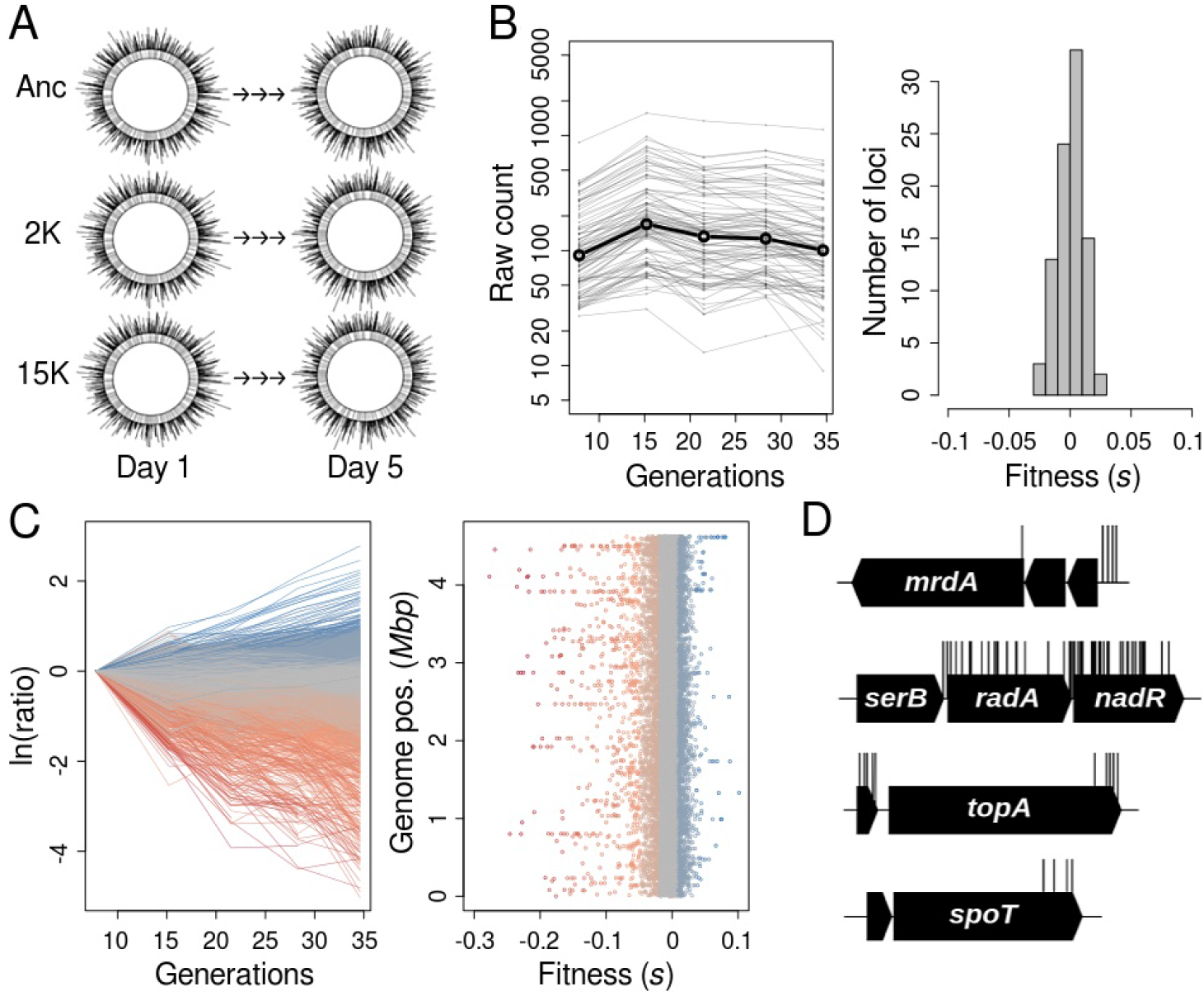
Overview of experimental procedures. **(A)** Several saturated, genome-wide insertion libraries in the ancestor and two evolved isolates (2K and 15K generations) from each of two LTEE populations (Ara+2 and Ara–1) were subjected to bulk competition and sequencing. **(B)** The abundance trajectories of well-known neutral loci were used to normalize coverage depth across time points, providing an internal reference to estimate selection coefficients (left). The values for this set of neutral loci were closely centered around zero (right). **(C)** Frequency trajectories of the whole mutant library in the ancestor (left), and mapping of the selection coefficient estimates along the chromosome (right). Colors indicate fitness effects, from deleterious (red) to beneficial (blue). **(D)** Examples of important sub-genic structure for known targets of selection including polar effects involving the preceding regions of the same transcription unit (*mrdA, nadR*, and *topA*), and tolerance to insertions in the C-terminal portion of essential genes (*serB, topA*, and *spoT*).

### Changes in the size and shape of the beneficial tail of the DFE

Using this approach, we first sought to characterize how the macroscopic shape of the DFE changed as fitness increased during the LTEE. The rate of fitness increase declined, such that half of the ∼70% gain typically seen at 50,000 generations had already occurred by 5,000 generations (*18*). We decided therefore to create transposon libraries in three genetic backgrounds: the ancestor (which we call “Anc”) and clones sampled from population Ara+2 at 2,000 (“2K”) and 15,000 (“15K”) generations, when fitness had increased by ∼25% and ∼50%, respectively. Despite these large fitness gains, Figure 2 shows that the overall shape of the DFEs remained similar, with one critical difference—namely, the fraction of beneficial insertion mutations is substantially larger in the ancestor than in the evolved backgrounds (6.8% for Anc versus 3.5% and 2.6% for 2K and 15K, respectively; *P <* 0.043 both cases, two-sample Kolmogorov–Smirnov [K-S] test). In contrast, the deleterious fraction is essentially constant across the three backgrounds (20.4% for Anc versus 17.2% and 16.1% for 2K and 15K, respectively; *P >* 0.083 both cases, two-sample K-S test). These patterns are consistent with analyses performed at the level of individual genes for both beneficial and deleterious mutations (Fig. 3, a-b).

**Fig. 2.**
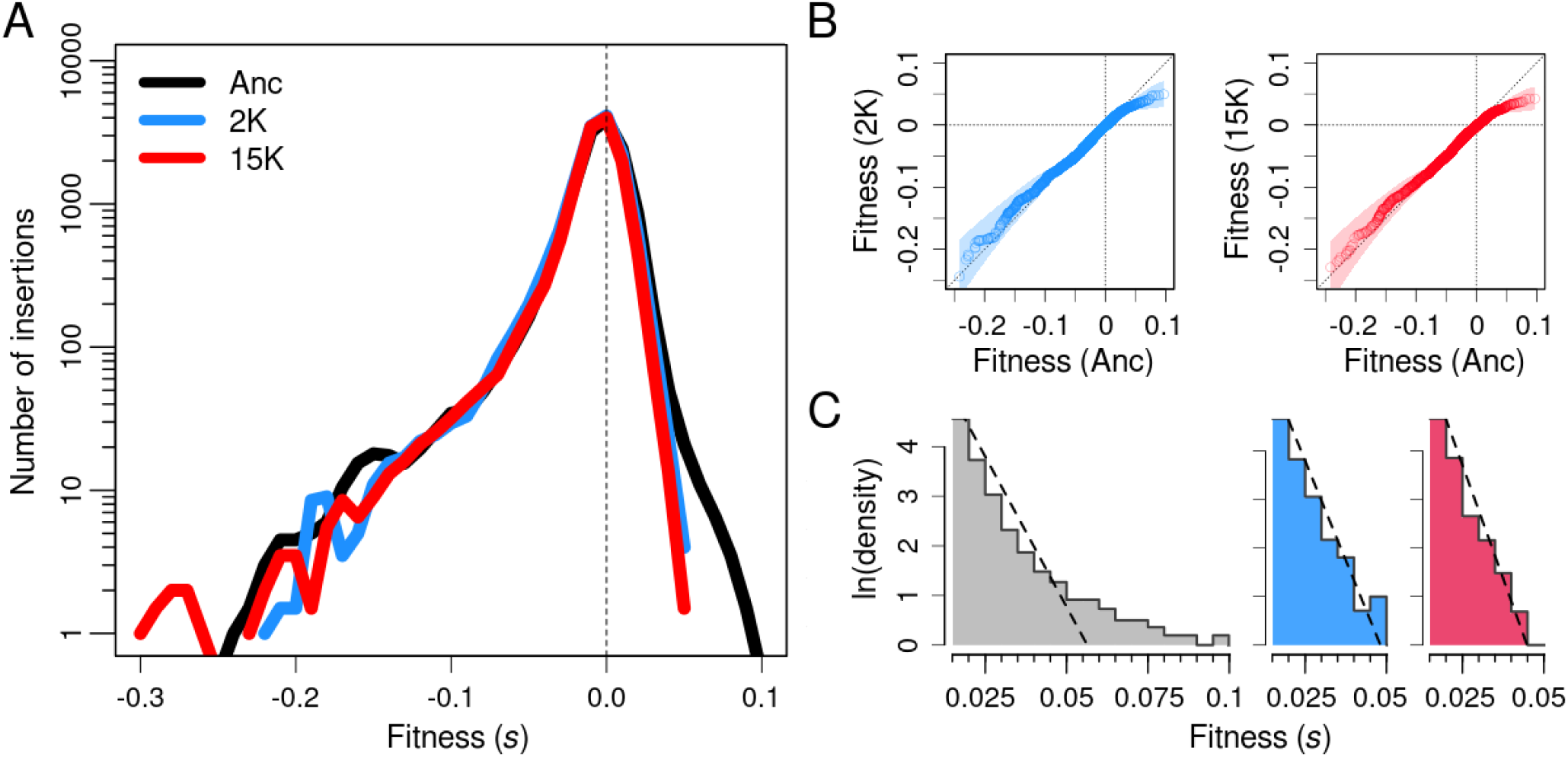
Change in the DFE along a large fitness gradient. **(A)** DFEs in the ancestor (black), 2K (blue) and 15K (red) evolved strains from population Ara+2. Note that the logarithmic scaling of the y-axis exaggerates minor, nonsignificant differences in the extreme deleterious tails. **(B)** Deleterious tails were unchanged during adaptation, as indicated by comparing the cumulative fitness distributions for the ancestor and 2K evolved strain (left), and for the ancestor and 15K strain (right). Shaded areas show 90% bootstrapped confidence intervals. **(C)** Beneficial tails were rapidly truncated during the LTEE, and they became exponentially distributed. Histograms show the best fits to exponential distributions (dashed lines) in the ancestor (gray), 2K (blue), and 15K (red) backgrounds. Note that all three x-axes use the same scale.

**Fig. 3.**
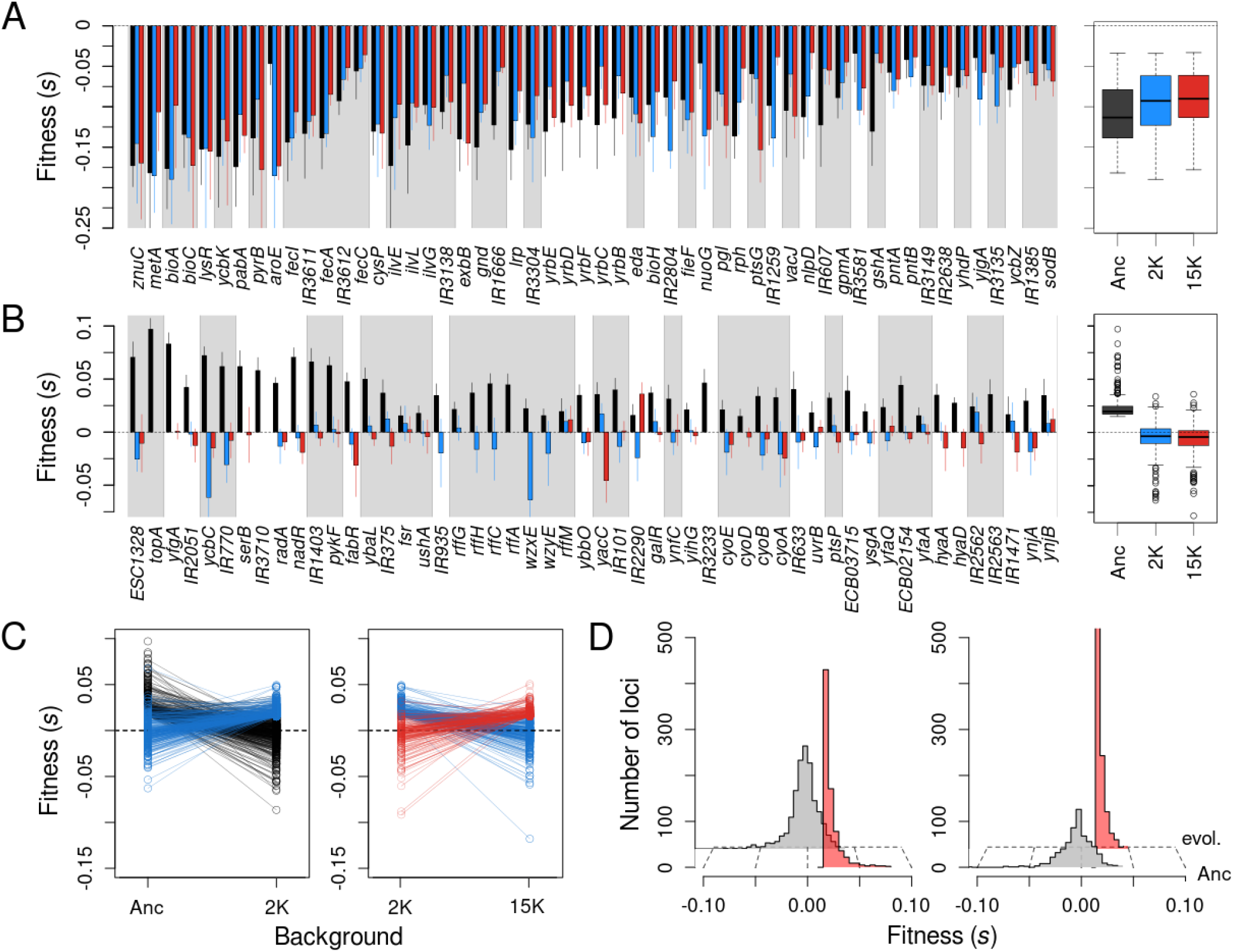
Effect of genetic background on the fitness effects of mutations in specific gene targets. **(A)** The genes and intergenic regions subject to the most severely deleterious mutants in three backgrounds in the Ara+2 lineage. Colors indicate ancestor (black), 2K (blue), and 15K (red) evolved strains. Gray shaded areas indicate loci in the same transcription unit. Values show the average across the different segments of each locus. Error bars indicate 90% confidence intervals. **(B)** The genes and intergenic regions with the most beneficial alleles in the ancestral background, and their fitness effects in the 2K (blue) and 15K (red) backgrounds. **(C)** Most of the beneficial mutations available to the ancestor became neutral or deleterious in the 2K background, while most beneficial mutations available in the 2K background were neutral or deleterious in the ancestor (left). The same general pattern occurs when comparing beneficial mutations in the 2K and 15K backgrounds (right). **(D)** More than 90% of initially beneficial mutations became neutral or deleterious in later generations (left). Likewise, more than 90% of beneficial mutations from later generations were neutral or deleterious in the ancestor.

We also examined whether these results depended on the particular evolutionary lineage that we chose to study. To that end, we measured the DFEs for clones sampled at 2,000 and 15,000 generations from population Ara–1, an independent lineage that accumulated a different set of beneficial mutations along its adaptive trajectory (Methods, Table S1). At least two major features distinguish the evolutionary history of this lineage from that of Ara+2. First, Ara–1 fixed a mutation in *topA* early in the LTEE, which confers the highest fitness benefit seen in this system for any single substitution (*19*). Mutations in this locus fixed in five of the twelve LTEE populations, but they never reached detectable frequency in Ara+2. Second, Ara–1 evolved a mutator phenotype, whereas Ara+2 retained the low ancestral mutation rate throughout the experiment; however, Ara–1 became hypermutable only after ∼21,000 generations, and hence poses no added technical complications to our study. Despite their independent histories, we obtained strikingly similar results for the two lineages, at both the macroscopic and microscopic levels (Fig. S3).

How do our findings compare to previous studies and expectations? An influential prediction based on statistical arguments is that the effects of beneficial mutations should be exponentially distributed when a population is well-adapted to its environment (*1, 12*). Despite some empirical support (*24*–*26*), the evidence remains inconclusive owing to a severe limitation of most studies: without detailed knowledge of a population’s evolutionary history, it is difficult to characterize its level of adaptation to a particular environment. Our data, by contrast, provides a uniquely powerful test of these ideas. We find that, indeed, beneficial mutations in the evolved backgrounds are well fit by an exponential distribution, whereas this distribution can be rejected for the ancestor (*P <* 0.001 for Anc versus *P =* 0.554 and *P =* 0.852 for Ara+2 clones 2K and 15K, respectively; one-sample K-S test). We also considered other alternative distributions, but the exponential provides the best fit for the evolved backgrounds (Methods, Table S2). Note that the exponential distribution is a special case of both the Weibull and gamma distributions, so it is not surprising that the data also fit well to them. These two distributions can be thought of as natural transitional shapes before reaching the limiting case of the exponential distribution. Indeed, the beneficial tail for the ancestor was fit to different degrees by both gamma and Weibull distributions (*P =* 0.035 and *P =* 0.29, respectively; one-sample K-S test), consistent with previous studies of viral and bacterial genotypes thought to be poorly adapted to their test environments (*26, 27*). Overall, our results support the view that, after an early period of rapid adaptation to a new environment, the distribution of beneficial mutations becomes exponential. Thus, by analyzing changes in the DFE in a temporal series of genetic backgrounds becoming better adapted to their environment, we can reconcile otherwise disparate pieces of evidence and provide insights relevant to many models of adaptation.

### Constancy of the deleterious tail of the DFE

The constancy of the deleterious tail we observe over time stands in contrast to a study that measured the DFE for 710 insertion mutations in hybrid yeast genotypes with fitness values spanning ∼20%, in which deleterious effects were significantly worse in the more-fit backgrounds (*28*). A potentially important difference is that the fitness variation among the yeast backgrounds was generated by crossing two distantly related strains, whereas we use a series of backgrounds from lineages undergoing adaptation to the same environment in which we assess the fitness effects of the new mutations. As further support for our findings, a companion study focused on the effects of deleterious mutations found no systematic changes in those effects across all of the LTEE lineages over 50,000 generations (*29*). In any case, theoretical predictions about the tail of deleterious mutations differ substantially and have been guided mostly by plausibility arguments (*14, 15*), and so all of these studies should help refine current models by clarifying the assumptions and narrowing the range of parameters.

### Changing identity of beneficial mutations and sign epistasis

Having examined how the macroscopic structure of the DFE changed as the bacteria adapted to the LTEE environment, we next sought to understand how the macroscopic changes emerged from changes at the level of genes and mutations. Figure 3a shows that deleterious mutations typically exhibit only slight epistasis across the three focal genetic backgrounds of the Ara+2 lineage. That is, the magnitude of their harmful effects may vary, but the tendency is for mutations that are deleterious mutations in the ancestor to remain deleterious in the evolved backgrounds, consistent with the observed constancy of the deleterious tail (see Fig. S4 for more details). In stark contrast, beneficial mutations are dominated by strong, sign-epistatic interactions (Fig. 3b). Only 5.9% of the mutations beneficial in the ancestor are still beneficial at 2,000 generations, with most becoming effectively neutral (76.9%) and some deleterious (17.2%) (Fig. 3c, left panel). This pattern also holds in the reverse direction: most beneficial mutations at 2,000 generations are neutral (78.4%) or deleterious (19.3%) in the ancestor (Fig. 3c, left panel). Similar patterns occur when comparing how fitness effects changed between 2,000 and 15,000 generations (Fig. 3c, right panel). Intrigued by the transitory nature of beneficial effects, we asked whether the overall DFE of the initially beneficial mutations retains even a slightly positive tendency at the later time points. In fact, it does not. The DFE of mutations that were beneficial in the ancestor becomes indistinguishable from a random sample of the parent distribution (Fig. 3d, left panel), and the same holds for the reverse scenario (Fig. 3d, right panel) (*P >* 0.116 both cases; two-sample K-S test). This regression to the mean persists even when we account for measurement noise around neutrality (Fig. S2).

What might explain this turnover in the identity of the beneficial mutations? In a previous study, the first five mutations to fix in one LTEE population were shown to exhibit diminishing-returns epistasis, such that their benefits declined in magnitude as the background fitness increased (*19*). However, it was unlikely *a priori* that these five early mutations would show sign epistasis because they were chosen precisely because their combination was favored by natural selection (*30*). By contrast, another study analyzed the co-occurrence of fixed mutations across 115 lines of *E. coli* that had evolved under thermal stress, and it found that sign epistasis was indeed common (*31*). Moreover, that study found that the prevalence of different types of epistasis reflected the modular architecture of cellular traits: mutations affecting different modules tend to interact more or less additively, while mutations impacting the same module tend to be redundant. We therefore investigated the extent of modularity in our data, and we found that beneficial mutations tend to cluster together in operons (Methods, *P <* 0.01). Mutations in the same operon typically alter the same cellular process in similar ways, and therefore the potential for redundancy at this functional level provides a simple explanation for why large sets of beneficial mutations disappear, and other sets emerge, as adaptation proceeds. Even without considering these specific details, the increased prevalence of sign epistasis with adaptation has also been predicted from general properties of the genotype-to-fitness map (*32*).

We identified a large set of loci that can produce beneficial mutations, including some known targets for adaptation in the LTEE (e.g., *topA, pykF, nadR*) (*20*). However, the fate of beneficial mutations is determined not only by their individual fitness effects, but also by their occurrence rate and the nature and intensity of their interactions with other beneficial mutations (*16, 17*). As a consequence, only a fraction of all possible beneficial mutations will contribute to adaptation in an evolving population. To gain further insight into this issue, we compared our data with metagenomic data previously obtained by sequencing whole-population samples from the 12 LTEE populations over the course of 60,000 generations (*21*). We see a significant, but fairly weak, correlation between our fitness estimates for mutations in the ancestor and the abundance of corresponding alleles during the LTEE (*r* = 0.27, Fig. 4a), and this correlation largely disappears when using the beneficial effects estimated in the evolved backgrounds. An important factor contributing to these weak correlations may be that our methods involve insertion mutations, which usually, but not always, cause losses of function (Fig. 1). While losses of unused functions have contributed to adaptation in the LTEE (*20, 33*), subtle changes that typically require point mutations have also been important in refining some functions (*16, 20, 34*). In contrast, the abundance of alleles in the metagenomic data correlates more strongly with the target size of the locus (*r* = 0.71, fig. 4b) (Methods). These patterns are consistent with intense competition among independently segregating beneficial mutations (*i*.*e*., clonal interference), a pervasive phenomenon in the LTEE (*21, 35*). Under intense clonal interference, the probability that particular beneficial mutations occur may shape genomic evolution even more than their individual fitness effects (*36*). In any case, the best linear model includes target size as the most explanatory single variable, but it also includes significant contributions from the fitness effects in both the ancestral and 2,000-generation genetic backgrounds (Fig. 4c, Table S3).

**Fig. 4.**
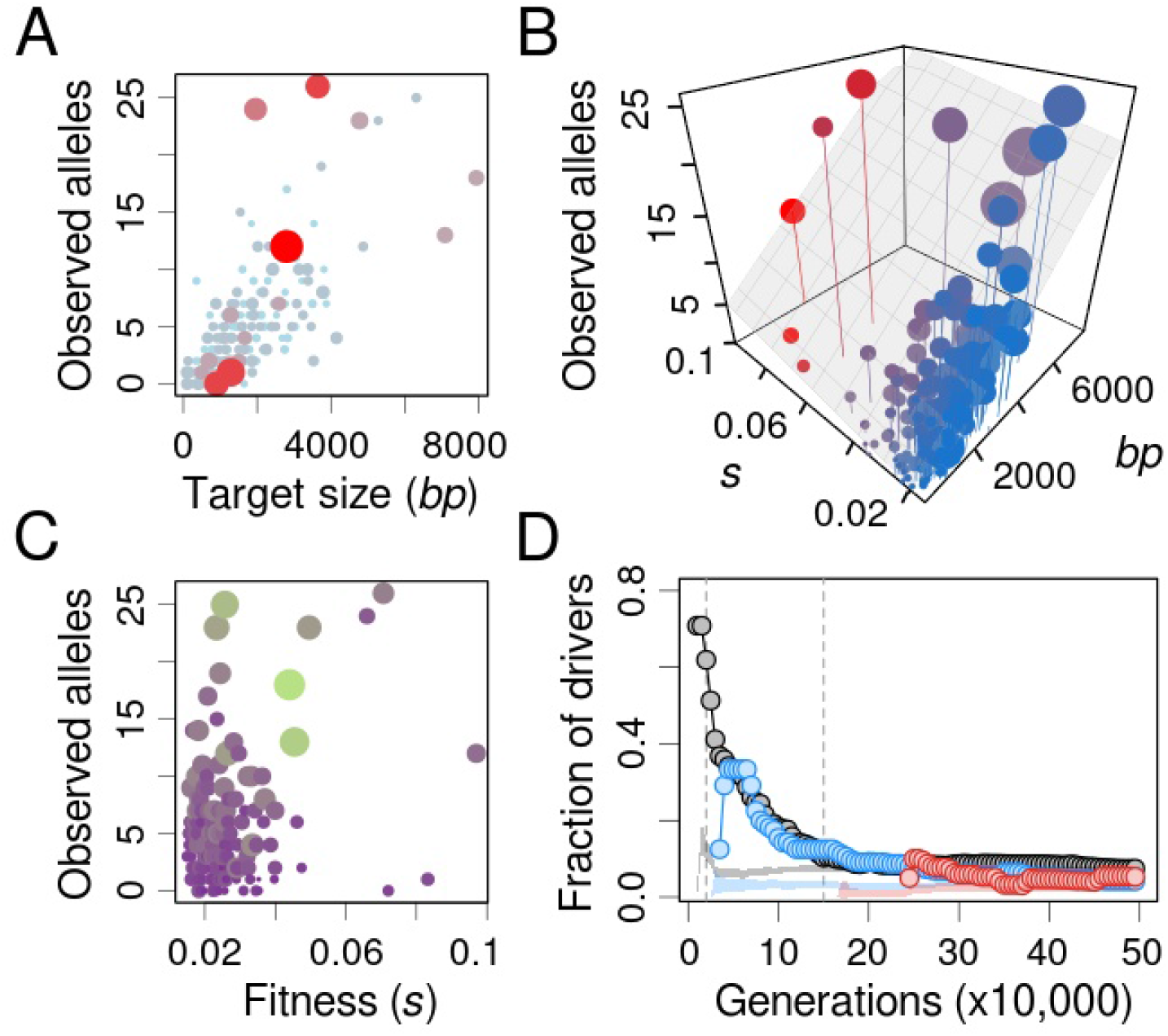
Determinants of evolutionary outcomes. **(A, C)** The prevalence of mutations in the LTEE is better explained by mutational target size (A, area and color of dots represent fitness) than by beneficial fitness effects measured in the ancestor (C, area and color of dots represent target size). **(B)** The best linear model for mutation prevalence includes fitness but is more strongly dependent on the mutational target size (area of dots represents target size, and color represents fitness). **(D)** The predictive capacity of DFEs as a function of time in the LTEE. Values show the fraction of numerically dominant alleles at each generation that were captured by the DFE measured in the ancestor (black), 2K (blue), and 15K (red) evolved strains. For the ancestor, we measured this fraction across all 12 LTEE populations; for the evolved backgrounds, the fraction includes only the focal population. Shaded areas show the null expectations based on randomly sampling neutral and deleterious mutations.

### Predicting future beneficial mutations as adaptation proceeds

Finally, given that sign epistasis is widespread, it is natural to ask for how long the information about the particular loci in the beneficial tail of a DFE can successfully predict the subsequent steps of adaptation. To address this question, we used the metagenomic data to record the alleles that were nearing fixation through time, and we calculated how many of those alleles corresponded to loci for which we detected beneficial effects. We found that the ancestral DFE predicted most of the loci where mutations became dominant early in the LTEE populations; the predictive power decays rapidly, but it was still evident for ∼15,000 generations (Fig. 4d). This decay was largely driven by lineages that evolved hypermutability early in the LTEE; when these mutator populations are removed from the analysis, the ancestral DFE retained significant predictive power through 50,000 generations (Fig. S5, a). In turn, the DFEs measured in the evolved backgrounds had less predictive power, and it took longer for their predictions to materialize; the latter effect may reflect the declining rate of adaptation. These patterns are reminiscent of work showing that parallel genomic evolution was more common early in the LTEE than in later generations (*20, 37*).

Why does the ancestral DFE have such predictive power, when it is estimated from insertion mutations that represent only a limited set of all possible mutations from a functional standpoint? To address this question, we quantified how many loci with frequent beneficial mutations in the LTEE include mutations with presumed loss-of-function effects. To that end, we assumed that nonsense, frameshift and structural variants cause losses of function. We find that these presumptive inactivating mutations contribute most of the initial adaptive mutations in the LTEE, and they represent a sizable fraction over the long run (Fig. S5, b). This result provides support for the “coupon-collecting” model of rapid evolution (*21, 31*), in which “rough-and-ready” loss-of-function mutations dominate the early phase of adaptation to a new environment owing simply to their high rates of occurrence. Under this model, many initially beneficial mutations are also redundant because they inactivate the same functional module. As a result, the model implies that fitness effects alone are a poor predictor of adaptive fixations, but taking target size into account can compensate for this uncertainty. This interpretation satisfactorily explains our findings that the initial drivers of adaptation are predictable despite widespread and strong epistasis, and that target size is the best predictor of beneficial alleles that fix early when a population encounters a new environment.

Taken together, our results shed new light on which aspects of adaptation to a novel environment are predictable, and which are not, given our current understanding of genetics and evolution. While the first steps of adaptation are not yet well-described by theory, we have shown that the identity of many drivers of early adaptation can be predicted from high-throughput empirical fitness data. By contrast, the long-term dynamics of adaptation are well-described using simple statistical arguments. However, predicting the genetic identity of the late drivers of adaptation is more difficult, and it will remain a challenge until a general theory of epistasis has been developed.

## Supporting information

Supplementary information includes: Materials and Methods, Figures S1 to S9, Tables S1 to S4 and Supplementary References.

## Acknowledgements

We thank Michael Baym and Anurag Limdi for valuable discussions; and Adrien Launay, Alexandra Baron, Romain Fernandes and Damien Roux for technical assistance. Funding: this work was supported by the European Commission under the 7th Framework Program (ERC Grant 310944 to O.T.), *Foundation pour le Recherche Médicale* (EQU201903007848 to O.T.), *Agence Nationale pour la Recherche* ANR GeWiEp (ANR-18-CE35-0005-0 to O.T.) and under the Horizon 2020 Framework Programme (MSCA-IF 750129 to A.C.). The work was also partly supported by the *Ministerio de Ciencia, Innovación y Universidades* (PID2019-110992GA-I00 to A.C.), a *Comunidad de Madrid “Talento”* Fellowship (2019-T1/BIO-12882 to A.C.), the US National Science Foundation (DEB-1951307 to R.E.L), and Michigan State University. Competing interests: We declare no competing interests.

