## Supplementary information includes: Materials and Methods, Figures S1 to S9, Tables S1 to S4 and Supplementary References. for "Predictability shifts from local to global rules during bacterial adaptation"

**This PDF file includes:**

Materials and Methods
Figures S1 to S9
Tables S1 to S4
Supplementary References

### Materials and Methods

#### Generation of transposon mutant libraries

We used the INSeq methodology (*1*) to create transposon mutant libraries in the ancestor and several evolved clones from the *E. coli* long-term evolution experiment (LTEE) (Table S1). We chose two populations, called Ara+2 and Ara–1, because they neither evolved hypermutability nor diversified into stably coexisting lineages during the first 15,000 generations. Many of the other LTEE populations evolved one or both of these features, which would complicate testing the hypotheses that we examine (ref. 26 in the main text). We employed a kanamycin-resistant version of the original INSeq transposon carried by the pSAM plasmid. The ends of this transposon encode recognition sequences for the restriction enzyme *Mme*I, which cuts 20 bp away from its binding site and thus allows the capture of the 14 bp adjacent to the insertion site (Fig. S1). The pSAM plasmid also carries the Himar1C9 transposase, an ampicillin-resistance marker, and RP4-oriT/oriR6K, a mobilizable and *pir*-dependent origin of replication. We maintained the plasmid in a *pir*-gene encoding *E. coli* donor strain, MG1655 MFD*pir* (*2*); from there, it was transferred to the *E. coli* recipient strains by standard agar-plate conjugation methods. Mating time was limited to 3-4 h, after which we selected transconjugants by spreading the cells onto LB agar plates supplemented with streptomycin (100 mg/L) and kanamycin (100 mg/L). Libraries were assembled from at least 100,000 colonies, which were obtained from at least 10 independent conjugation mixtures. The colonies were scraped off the plates, pooled in saline solution, and glycerol was added (10% final) for storage at –80°C.

#### Bulk competition experiments

We propagated aliquots of the mutant libraries in the same conditions as those used in the LTEE: 1/100 daily dilutions in 50-mL flasks filled with 10 mL of DM25 and incubated at 37°C in an orbital shaker (*3*). To ensure that the initial cell density was similar to that used in the LTEE (~5 x $10^7$ cells/mL), we inoculated several flasks for each experiment with a different aliquot from the thawed libraries, using only those with appropriate densities based on viable cell counts (and discarding the others). The competition experiments ran for 5 or 8 days (Ara+2 and Ara–1 samples, respectively); we stored the remaining culture after each serial passage at –80°C for later analyses. We monitored the populations based on viable cell counts to verify that they grew for the expected ~6.6 generations each day.

#### Generation of DNA libraries for sequencing

The thawed stocks from the bulk competition experiments were pelleted and subjected to genomic DNA extraction using the DNeasy Blood & Tissue kit (Qiagen). Owing to the relatively low cell density supported by DM25, the purified DNA was subjected to high-fidelity, unbiased whole-genome amplification using the *Phi29*-polymerase-based REPLI-g midi kit (Qiagen). The resulting DNA was digested by *Mme*I (NEB), concentrated using a vacuum concentrator, and run on a 1% agarose gel. The expected size of the transposon plus the adjacent genomic DNA is ~1.5 Kb, so we excised from the gel the corresponding region, using as a reference the 1.2-1.5 Kb bands from the Quick-Load 100 bp DNA ladder (NEB). To enrich the excised samples for transposons, the INSeq method takes advantage of the fact that *Mme*I digestion results in dinucleotide, sticky-end fragments; custom-made PCR adaptors could then be added in a simple ligation reaction (Fig. S1, Table S4). We performed the ligations using T4 DNA ligase (NEB). Ligation products contain the 14 bp adjacent to the insertion site flanked by the transposon and the custom-made adaptor. Using these known sequences for annealing, we used a 28-cycle PCR to amplify a small region containing the 14 bp of interest (Fig. S1, Table S4). All PCR reactions were carried out using the HiFi DNA Polymerase (KAPA Biosystems). The resulting molecules (116 bp and 148 bp in size for the Ara+2 and Ara–1 samples, respectively) were isolated by electrophoresis and band excision on a 2% agarose gel. To add the required indices for multiplexed, paired-end sequencing, we subjected the samples to a secondary, 18-cycle PCR using the Nextera XT DNA Library Prep Kit (Illumina). The final Nextera libraries were purified, quantified, and sent for sequencing on Illumina platforms. The Ara+2 libraries were sequenced using a HiSeq platform (Integragen, Evry, France), while the Ara–1 libraries were sequenced using a MiSeq platform (Bichat Hospital, Paris, France). After filtering (see below), the libraries had on average ~3.55 and ~0.39 million reads, mapping to an average of ~0.33 and ~0.27 million different insertion sites for Ara+2 and Ara–1, respectively. To control for potential technical biases, libraries for the ancestor were processed using both transposon capture and sequencing methods (Fig. S6).

**Insertion location, filtering, and abundance determination**

We developed a series of custom scripts to map transposon insertion sites, discard low-quality cases, and quantify the abundance of each insertion allele. We first used a Python script to extract the captured 14-bp genomic fragments from the Illumina sequence reads. This step was accomplished using the module "regex" to identify any of the constant sequences expected to flank the captured fragments, up to a maximum edit distance of 1 (*i.e.*, one position indel or mismatch) (Fig. S1). Second, we used the BWA algorithm (Li et al., 2009) to map these genomic fragments to the *E. coli* B str. REL606 reference genome (NCBI reference: NC_012967), again with a maximum edit distance of 1. We only retained fragments for which the best hit mapped to a single genomic location, thus excluding repeated regions from the analysis. Pairs from paired-end reads were merged except when they gave discordant results (in these cases, only the one with the best alignment was retained). These filtered read mappings were then compiled in a list with all the unique insertion sites, the number of reads per site and the identity of the corresponding locus. As loci, we considered both open reading frames and intergenic regions, but we excluded repetitive and mobile genetic elements. Finally, to characterize sub-genic effects (Fig. 1, D), we divided each locus into equally sized segments (5 for the Ara+2 dataset, and 3 for the lower-coverage Ara–1 dataset; in both cases, loci were not divided when <100 bp in size), and we calculated the abundance of all insertions that mapped onto the same segment. In preliminary tests we observed that excluding insertion sites with very low counts reduced the number of artifacts in downstream analyses. This effect presumably occurs because very low abundance insertions are enriched for PCR and Illumina sequencing artifacts. Therefore, for each segment, we pooled together only insertions with

>10 or >3 total counts across all timepoints (Ara+2 and Ara–1, respectively). All the following analyses were conducted using the data for these pooled segments.

**Fitness estimates and outlier removal**

To estimate the fitness of the pooled insertion mutants for each gene or sub-genic region, we calculated the slope of their frequency trajectories relative to that of a set of presumed neutral loci. The natural logarithm of this slope approximates the selection coefficient from classical population genetics, and it is related to the ratio of realized growth rates (the standard fitness metric used in the LTEE literature) by a scaling factor of ln 2 (Fig. S7, a) (*4*). The set of presumed neutral loci includes genes annotated as cryptic and the L-arabinose transport and catabolism operons (L-arabinose auxotrophy has been widely used as a neutral marker in the LTEE). In addition, we only considered as neutral their internal segments, in order to avoid spurious deviations from neutrality associated with polar or change-of-function effects (e.g., Fig. 1D). We estimated slopes by fitting a linear regression model to the data using the lm() function in R. We weighted the regression using the counts at each time point (using the "weight" argument), in order to reduce the noise associated with low counts. We found this weighting improved estimates in the deleterious range (Fig. S7, b). Mutants with fewer than 3 non-zero timepoints were discarded.

In tests with the neutral set, we observed that most poor-quality fits could be filtered out simply by excluding mutants with very low initial abundance (cutoffs for exclusion of <10 or <3 counts at the first timepoint for Ara+2 and Ara–1, respectively). However, when the much larger complete set of loci was analyzed, we noticed several instances in which marked outliers were characterized by increasingly steep slopes at later timepoints. These cases probably reflect hitchhiking of individual insertion alleles with *de novo* beneficial mutations that arose during the competition experiments. Consistent with this interpretation, these outliers were proportionately more common in the higher-coverage Ara+2 dataset, presumably because of the increased resolution and hence ability to detect these rare events. To remove these outliers, we filtered them out using the standard error of the regression (cutoffs of 0.01 and 0.015 for Ara+2 and Ara–1, respectively). We also noticed a few highly beneficial mutations in the ancestor from the Ara+2 dataset that we suspect constituted another class of more subtle artifacts. Our suspicion arose from two facts: first, they were completely absent from the ancestor data for Ara–1; second, their fitness markedly deviated from the fitness measured for the other insertions in the same locus. One explanation for these artifacts is that they arose from some type of PCR bias. However, MmeI digestion ensures that genomic fragments are of uniform length (1.5 Kb), thus preventing any significant size-related extension rate differences across fragments (*5*). Other types of PCR bias cannot be excluded, but they probably play a minor role, as shown in a recent study comparing PCR-free versus PCR-based transposon enrichment methods (*6*).

Alternatively, we think the likely explanation is that these artifacts represent even rarer hitchhiking events, ones in which an insertion occurred in a genetic background harboring either a preexisting beneficial mutation or a secondary beneficial insertion. In such cases, the insertion would behave as a beneficial mutation over the entire time course, and thus it cannot be identified and discarded based on a high standard error of the regression. These rare events could even make deleterious mutations appear neutral or beneficial. To address this issue, we systematically compared all segments for each locus, and we removed those internal segments that deviated by > 0.01 in fitness from the locus average (except in cases where the deviation represents < 25% of the locus average, to avoid being too strict with loci displaying large deleterious effects; see Fig. S2, b). We retained N- or C-terminal segments, on the grounds that such deviations may represent polar or change-of-function effects; however, to retain them when beneficial, we further required that other beneficial mutations were found elsewhere in the same operon.

Overall, after imposing all of these filters for the Ara+2 dataset, loci retained an average of 3.5 ± 1.3 internal segments (mean ± SD; Anc: 3.2 ± 1.3, 2K: 3.7 ± 1.3, 15K: 3.6 ± 1.3), with each internal segment having an average of 7.2 ± 5.8 independent insertion alleles (Anc: 7.8 ± 6.3, 2K: 7.6 ± 6.1, 15K: 6.1 ± 4.8). For the Ara–1 dataset, loci retained an average of 2.4 ± 0.8 internal segments (mean ± SD; Anc: 2.4 ± 0.8, 2K: 2.4 ± 0.8, 15K: 2.3 ± 0.8), with each internal segment having on average 10.6 ± 8 independent insertion alleles (Anc: 12.4 ± 9.4,, 2K: 10.2 ± 7.9, 15K: 9 ± 6.8).

### Statistical methods

We performed all statistical tests using either built-in functions or publicly available packages from the R programming language (version 3.6.3). To fit linear models, we used the built-in lm() function. We used the package “Matching” to implement a bootstrap version of the Kolmogorov-Smirnov test, which corrects for the fact that the empirical distributions of fitness effects are discrete in nature (*7*). We used the package “Fitdistrplus” to find which of the distributions commonly used in the literature provided the best fit to the beneficial tails (Table S2) (*8*). For all tests conducted on both the deleterious and beneficial tails, we included only those values outside the range of neutrality, as established from the DFE for the set of presumed neutral loci (Fig. S8). To assess whether measurement error might drive the observed results, we resampled the data using the logarithm of the regression standard errors as weights. Weighted resampling with replacement was conducted 200 times for each genetic background and tail of the distribution, and the resulting datasets were used to repeat the tests associated with Figures 2B and 2C. These analyses confirmed that our main results hold even when accounting for measurement error, both for the deleterious and beneficial tails (Fig. S9). We also used the resampled datasets to estimate 90% confidence intervals around the cumulative DFEs (Fig. 2B).

To test whether beneficial mutations in the ancestor tended to cluster together in operons more often than expected by chance, we sampled without replacement an equal number of loci from the reference genome and computed the frequency of having another member from the same transcription unit in the sample. The reported p-value is the proportion of cases in 100 simulations that show an equal or higher frequency of clustered loci than the observed data. Operons in which mutations typically have small fitness effects should be underrepresented because most mutations will fall below the threshold of neutrality. To avoid this potential bias, we focused these analyses on the loci that harbored the 50% most beneficial mutations observed. The list of transcriptional units for the ancestor was retrieved from the BioCyc database (biocyc.org/group?id=:ALL-TRANSCRIPTION-UNITS&orgid=ECOL413997, accessed on 2/23/2022).

#### DFE-based predictions versus allele prevalence in the LTEE

We obtained the metagenomic time-series data for the LTEE from https://github.com/benjaminhgood/; genomic positions were used to standardize annotated loci labels as needed. For every non-synonymous allele observed in the metagenomic data, we recorded the number of lineages in which a similar allele was detected, the maximum frequency it reached, and the timepoint at which this maximum was attained. By similar allele, we mean any other non-synonymous located in the same gene or sub-genic region. Six of the LTEE lineages evolved hypermutability at various generations, which introduced many "passenger" mutations (i.e., neutral or mildly deleterious mutations that reach high frequency due to their linkage with a beneficial "driver" mutation) (*9*). Such passengers are typically unique because they are drawn from a very large pool of potentially neutral or mildly deleterious mutations. Therefore, we restricted our analyses to similar alleles (as defined above) that arose in at least 2 independent lineages. In addition, to ensure that these alleles represented beneficial mutations, we only considered those alleles that reached a frequency of at least 50% at some point in the corresponding metagenomic trajectory. When attempting to correlate the abundance of these presumptive driver alleles with our fitness data, we realized that working at the sub-genic and even the locus level relied on sparse data and thus had low statistical power. To overcome this limitation, we performed the analyses at the operon level; fitness was computed as the maximum value recorded for any of its constituents (including sub-genic regions), and target size was calculated as the sum of relevant constituent lengths (i.e., excluding sub-genic regions and loci without beneficial effects). We followed the same approach when we looked at how well the DFEs predicted subsequent steps of adaptation. To compute the fraction of drivers that were captured by the DFE, we retrieved the alleles reaching ≥50% frequency at a given timepoint and counted how many of them belonged to operons present in the beneficial tail of the corresponding DFE. In the analyses with the ancestor, we required the allele to reach the 50% threshold in at least 2 LTEE lineages. In the same analyses, except performed with the 2K and 15K genetic backgrounds, we only considered alleles that surpassed the threshold in their own lineage (Ara+2 or Ara–1), discarding those alleles already present above this threshold at 2K or 15K generations, respectively. Note that despite the focus on a single lineage, the mutations still come from the filtered "driver" set; hence, lineage-specific singletons were not considered. To compute a null expectation for these analyses, we ran them on random samples from the neutral and deleterious fractions of each DFEs. The samples were drawn without replacement and with a size equal to the beneficial mutation set to which they were compared. The shaded areas in Figures 4 and S3 show the average of 50 simulations ± three times the standard error of the mean.

### Supplementary Figures

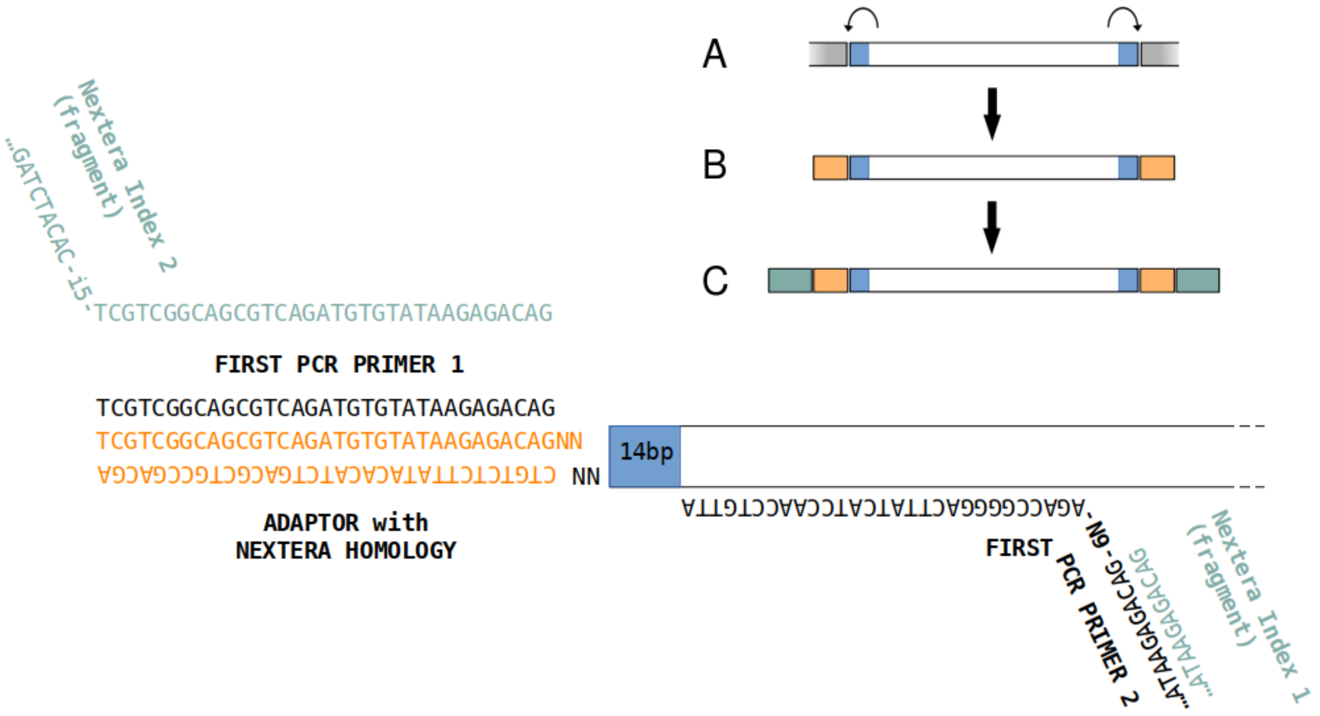


**Fig. S1. Procedure for capturing the transposon flanking regions. (A)** Genomic DNA is extracted in bulk, digested with *MmeI*, and separated by gel electrophoresis. **(B)** Transposon-sized fragments are ligated with double-stranded oligonucleotide adapters. **(C)** Two rounds of PCR are performed to add indices for Illumina sequencing.

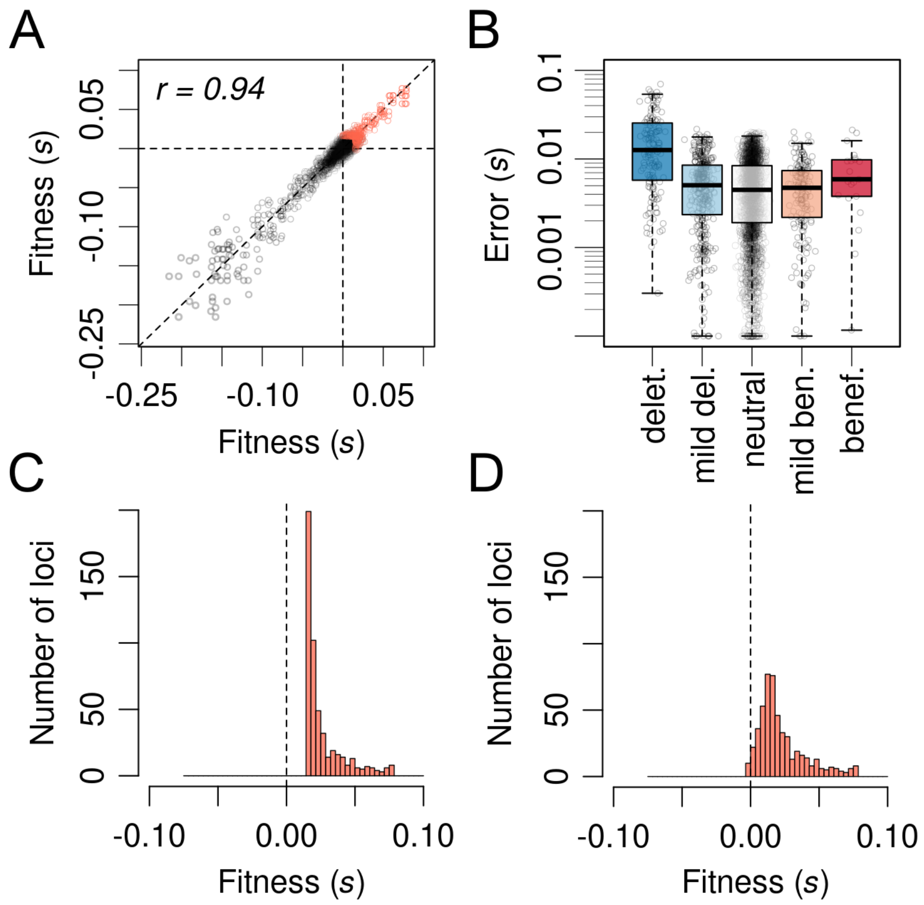


**Fig. S2. Reproducibility of fitness estimates (A)** Pairwise comparisons among segments from the same locus show high reproducibility. Pearson's coefficient, *r*, is in upper left corner. **(B)** Measurement error among pairwise replicates scales with the magnitude of fitness effects (deleterious and beneficial: $|s| > 0.05$; mild: $0.05 > |s| > 0.015$; neutral: $|s| < 0.015$. **(C, D)** Despite measurement noise, the shape of the DFE for beneficial mutations **(C)** retains a marked shift towards positive values when constructed using different segments from the same locus **(D)**.

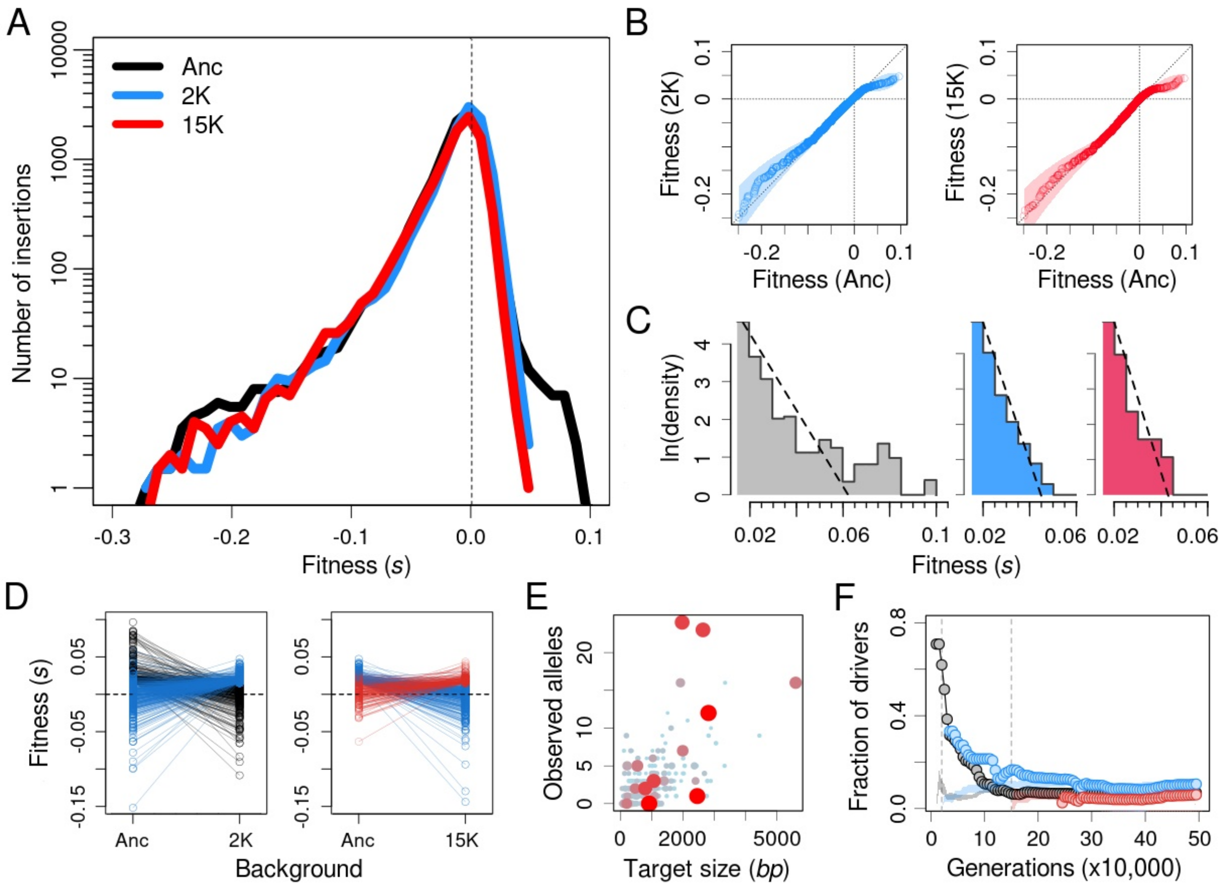


**Fig. S3. Change in the DFE along a second large fitness gradient. (A)** DFEs in the ancestor (black), 2K (blue) and 15K (red) evolved strains from population Ara–1. Note the logarithmic scaling of the y-axis. **(B)** Deleterious tails were unchanged during adaptation, as indicated by comparing the cumulative fitness distributions for the ancestor and the evolved strains (2K, left; 15K, right). Shaded areas show 90% bootstrapped confidence intervals. **(C)** Beneficial tails were rapidly truncated during the LTEE, and they became exponentially distributed. Histograms show the best fits to exponential distributions (dashed lines) in the ancestor (gray), 2K (blue) and 15K (red) backgrounds. Note that all three x-axes use the same scale. **(D)** Most beneficial mutations available to the ancestor become neutral or deleterious in the 2K background, while most beneficial mutations available in the 2K background were neutral or deleterious in the ancestor (left). The same general pattern occurs when comparing beneficial mutations in the 2K and 15K backgrounds (right). **(E)** The prevalence of mutations in the LTEE is correlated with the mutational target size (area and color of dots represent fitness). **(F)** The predictive capacity of the DFEs as a function of time in the LTEE. Values show the fraction of numerically dominant alleles at each generation captured by the DFE measured in the ancestor (black), 2K (blue), and 15K (red) evolved strains. Shaded areas show the null expectations based on randomly sampling neutral and deleterious mutations. The results shown here in all panels **(A-F)** for LTEE population Ara–1 are strikingly similar to those seen for the independently evolving Ara+2 population (see Fig. 2A-C, Fig. 3C, and Fig. 4A,D).

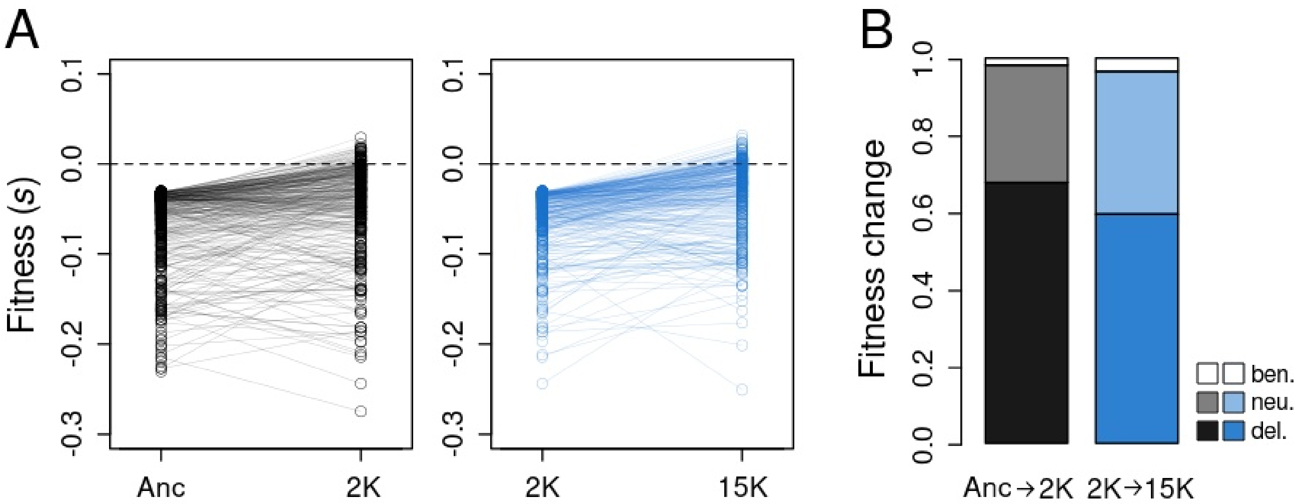


**Fig. S4. Patterns of epistasis among deleterious mutations in Ara+2 lineage. (A)** Most deleterious mutations in the ancestral background remain deleterious in the evolved 2K background (left), and most deleterious mutations in the 2K background remain deleterious in the 15K background (right). **(B)** The fraction of deleterious mutations with deleterious, approximately neutral, and beneficial effects in more fit backgrounds. Most changes from deleterious to neutral or beneficial effects occur in mutations with fitness effects close to the detection limit for neutrality in the earlier background ($s$ = –0.015).

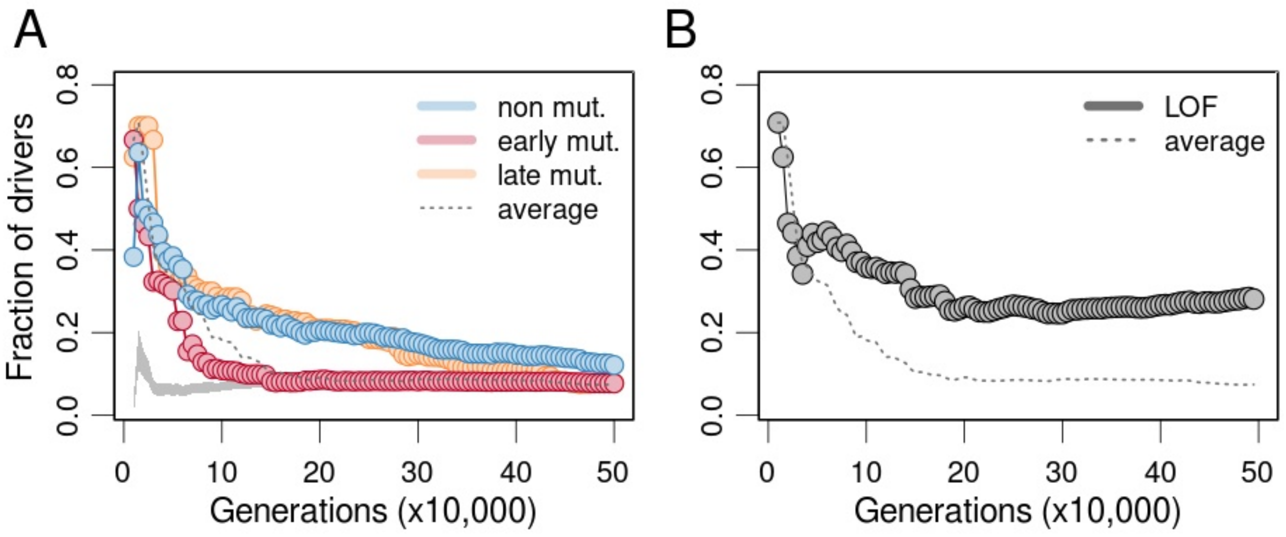


**Fig. S5. Loss-of-function mutations and the predictability of adaptation. (A)** Predictive capacity of the ancestral DFE as a function of time for different sets of LTEE lineages. Colors indicate analyses performed separately using data from the non-mutator (blue), early mutator (orange), and late mutator lineages (red). The dashed line shows the aggregate average when pooling all lineages, as shown in Fig. 4D. The shaded area shows the null expectation based on randomly sampling neutral and deleterious mutations. **(B)** The fraction of loci harboring high-frequency beneficial mutations that show mutations with presumed loss-of-function effects (nonsense, frameshift, and structural variants). The fraction of these important targets of adaptation captured by the DFE measured in the ancestor is shown for comparison (dashed line).

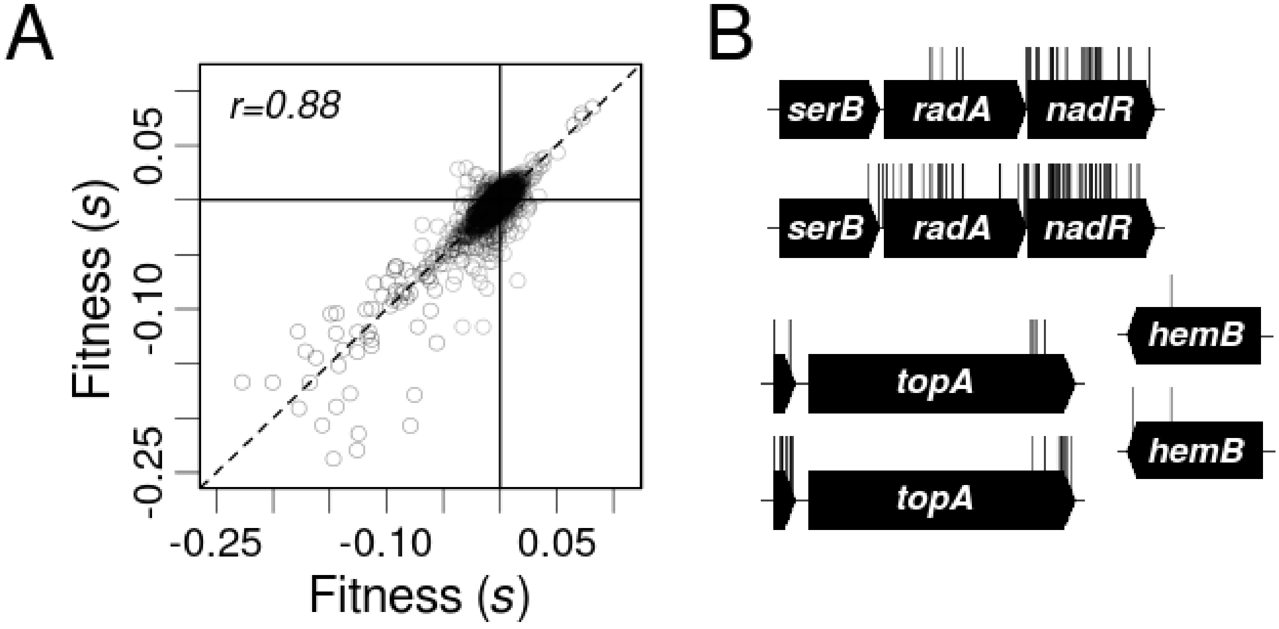


**Fig. S6. Comparison of fitness estimates between two datasets. (A)** Correlation between selection coefficients measured in the ancestral genetic background in independent datasets for two LTEE lineages (Ara+2 on the x-axis; Ara–1 on the y-axis). Pearson's coefficient, *r*, is shown in the upper left corner. **(B)** Lower coverage in the Ara–1 dataset resulted in lower resolution at the sub-genic level. This difference is best illustrated by mapping insertions in essential genes for which insertions in the C-terminal end are neutral (*hemB*) or even beneficial (*serB*, *topA*). The diversity of insertion sites in these loci is consistently lower in Ara–1 (top alignments) than in Ara+2 (bottom alignments).

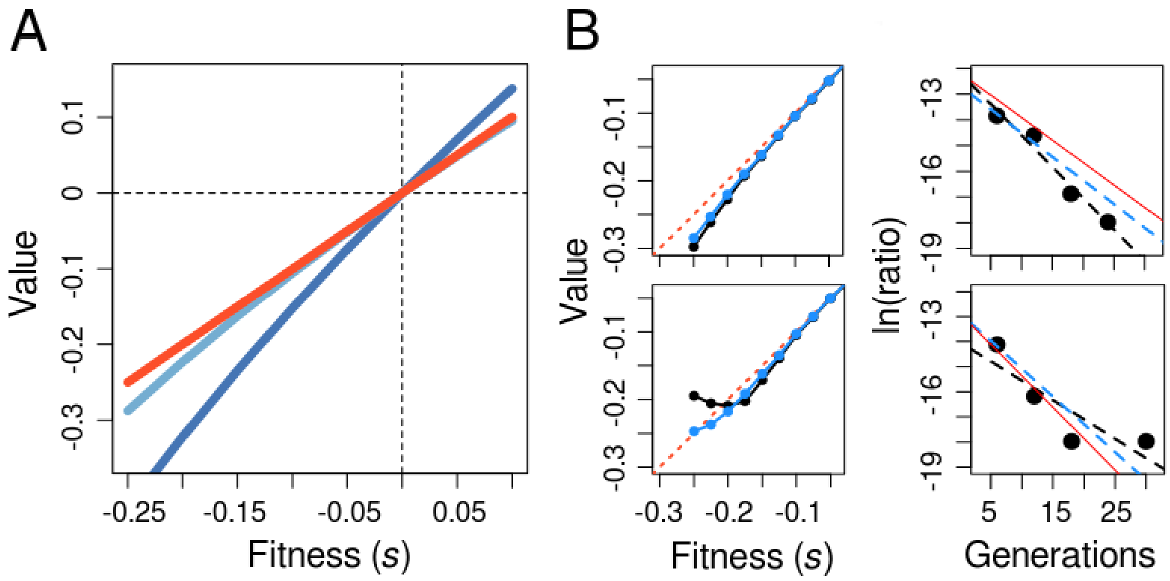


**Fig. S7. Measuring fitness effects from bulk competition data. (A)** Relationship between the fitness effect (red; often called the selection coefficient, *s*, in population-genetics literature) and the ratio of realized growth rates (dark blue; typically used as the measure of relative fitness, *W*, in the LTEE). Empirical estimates of *s* based on the slope of allele frequency trajectories make the approximation $s \approx \ln(1 + s)$, which equals *W* multiplied by ln(2) (light blue) (*4*). Note, however, that this approximation starts to deviate at large values of *s*. **(B)** Weighting the regression by the abundance of each allele improves estimation of *s*. This improvement is most important at low values of *s*, when the abundance of a highly deleterious allele approaches zero at later timepoints. Lines show the true *s* (red), standard regression estimate (black), and weighted regression estimate (blue). Top left panel shows results from a set of simulations with a single deleterious allele starting at a frequency of $7 \times 10^{-7}$, and the top right panel shows a single run with $s = -0.18$. Bottom left panel shows results from simulations in which the deleterious allele started at a frequency of $2 \times 10^{-7}$, and the bottom right panel shows a single run with $s = -0.25$. Each symbol in the two left panels shows the average of 100 simulations at each value of *s*. Simulation model adapted from (*10*), using a total population size of $6 \times 10^{7}$.

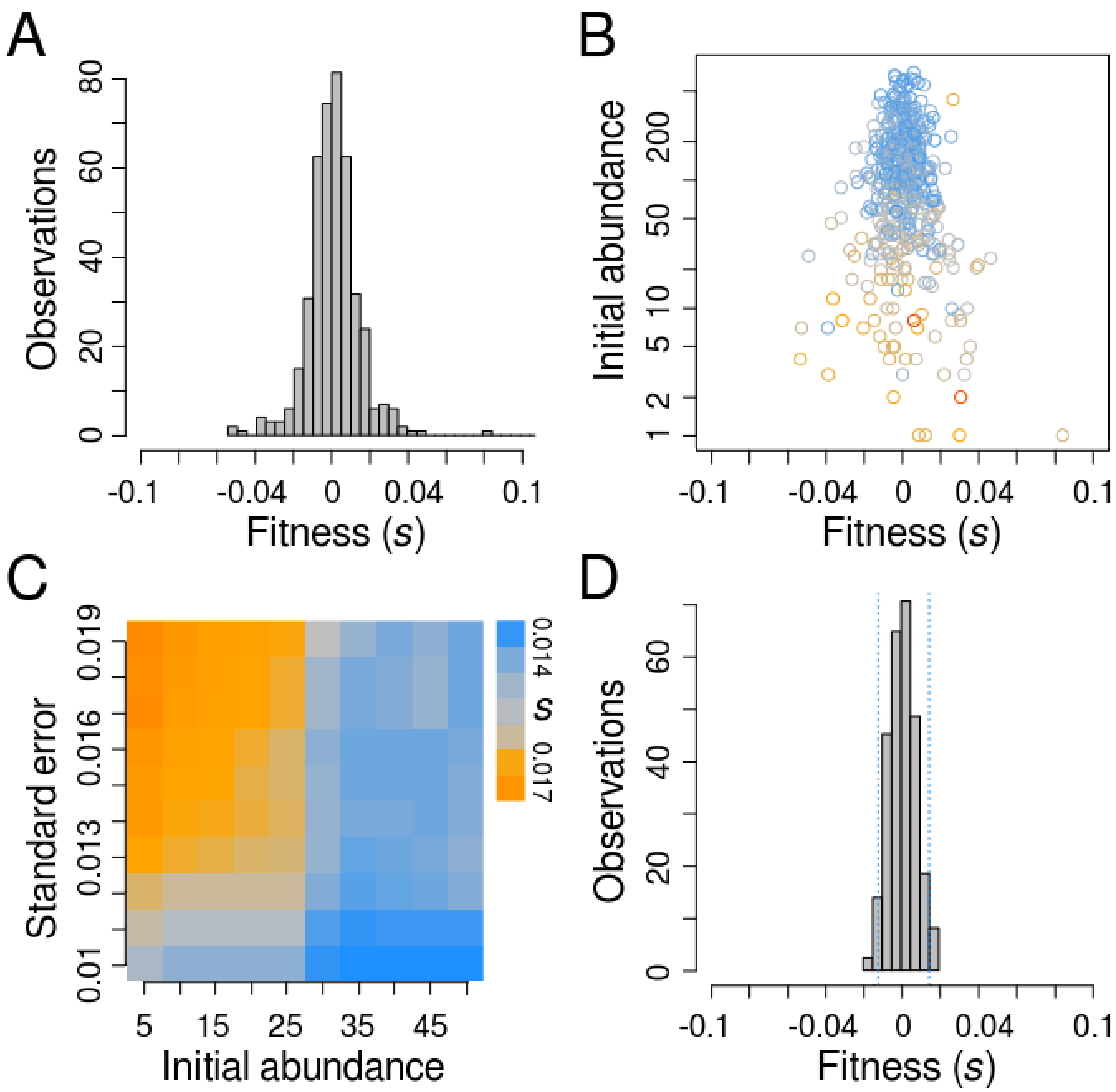


**Fig. S8. Data filtering and the threshold of neutrality. (A)** Raw distribution of fitness effects for presumed neutral loci in datasets from the ancestor and both evolved Ara+2 strains. **(B)** Most small deviations from zero can be explained by sampling error, while larger deviations result from low initial abundance (y-axis, note logarithmic scaling) or poor linear fit (colors from blue to gray to orange indicate increasing standard errors). **(C)** After filtering artifacts, we scanned combinations of initial abundance and standard error to identify selection coefficients encompassing 97% percent of the data (colors, see scale bar on right). **(D)** Final distribution of selection coefficients for presumed neutral loci after applying all the filters. The blue dotted lines indicate the resulting threshold for approximate neutrality (–0.015 < *s* < 0.015).

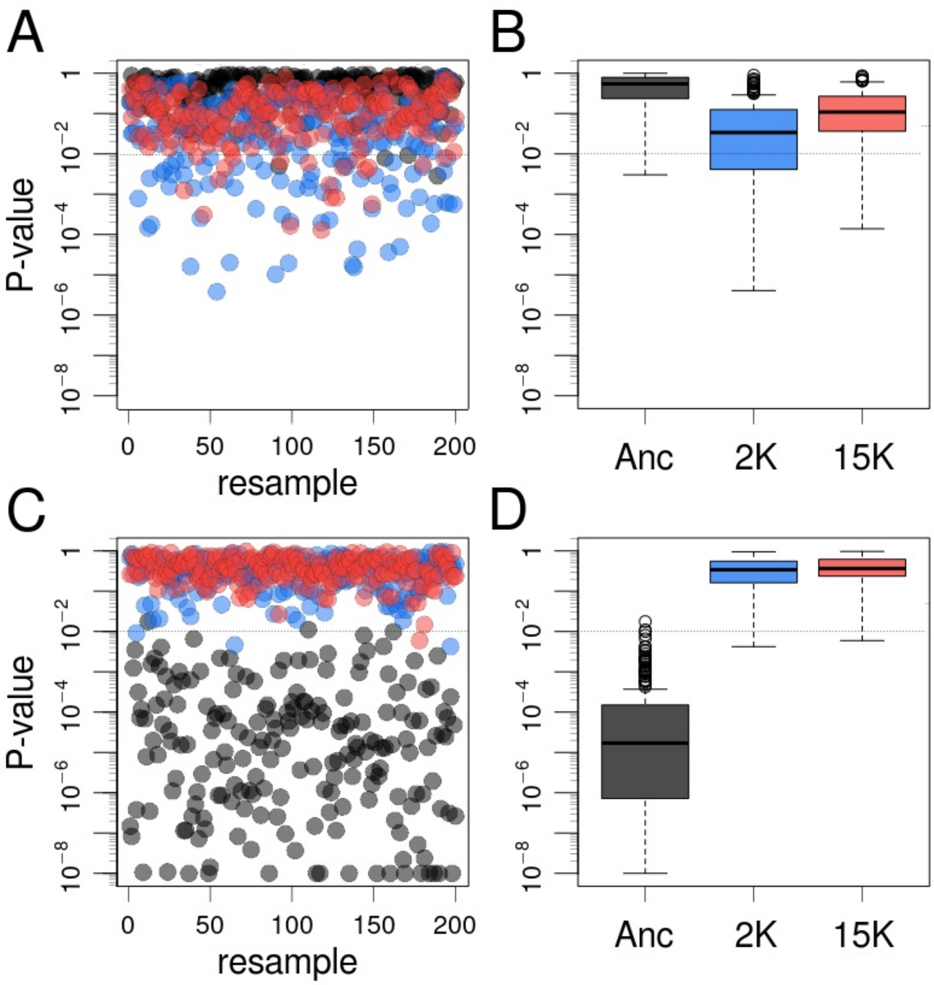


**Fig. S9. Measurement error and reliability of tests on the shape of DFEs.** To assess the role of noise in statistical analyses of the shape of DFEs, we resampled the distribution tails using regression standard errors as weights. **(A)** P-values from replicated two-sample Kolmogorov–Smirnov (K-S) tests in which the ancestor's deleterious tail was compared with 200 resampled deleterious tails obtained from the ancestor (used as control, black), 2K (blue), and 15K (red) backgrounds from the Ara+2 lineage. Note the logarithmic scaling of the y-axis. **(B)** Boxplots show that deleterious tails in the more-fit evolved backgrounds are similar to the ancestor's tail, although for the 2K background the results are ambiguous (average P-values were 0.51, 0.096 and 0.18 respectively). **(C)** P-values from replicated one-sample K-S tests in which an exponential fit was made for 200 resampled deleterious tails from the ancestor (black), 2K (blue), and 15K (red) backgrounds. **(D)** Boxplots show that the beneficial tails in the evolved backgrounds are well fit by an exponential distribution, but this distribution is rejected for the ancestor (average P-values are $1.7 \times 10^{-5}$, 0.34, and 0.37 respectively).

### Supplementary Tables

**Table S1. Strains used in this study**

| **Name** | **strain ID** | **mutated loci[1]** | **source** |
|---|---|---|---|
| Ancestor | REL606 | None | (*3*) |
| Ara+2 2K | REL1159A | *pykF, wcaM, infB, yhgI/gntT, pitA, spoT, rbsD–[rbsB], ECB_03710* | (*11*) |
| Ara+2 15K | REL7184A | *insL-2/lon, ybaL, [nmpC]–[insK-1], lrp, rpsA, appC, flgB, rnb, speG, pykF, [lpp]–[ydiM], [lpp]–[ydiM], [wzy]–[wcaM], wcaM, nupC/yfeA, purL, [serV]–insL-5, yqeB, [ECB_02797]–ECB_02822, mreB, fis/yhdJ, spoT, rbsD–[rbsA], ilvL, yihP, hslU, iclR, metH, sgcR–insA-27, nadR, arcA* | (*11*) |
| Ara-1 2K | REL1164A | *ECB_00736/ECB_00737, topA, pykF, spoT, glmU/atpC, rbsD–[yieO]* | (*11*) |
| Ara-1 15K | REL7177A | *mokC/nhaA, pcnB, insB-2, araJ, ybaL, [nmpC]–[ECB_00513], mrdA, nagC, ompF/asnS, dhaM, ldrC/chaA, narI/ECB_01206, topA, pykF, ynjI, yedW/yedX, [manB]–[cpsG], yegI, ECB_02816, yghJ, ebgR, infB, arcB, gltB, yhdG/fis, rpsM, malT, glpE, spoT, glmU/atpC, rbsD–[yieO], hslU, pflC, iclR, fimA, nadR* | (*11*) |

[1]Note: annotations follow conventions from https://barricklab.org/shiny/LTEE-Ecoli/

**Table S2. Fits of several common distributions to beneficial tails of DFEs.**

| **One-sample Kolmogorov-Smirnov test, p-values[1]** | | | | | | |
|---|---|---|---|---|---|---|
| Background | Exponential | Gamma | Weibull | Normal | LogNorm | Logistic |
| Anc | $<10^{-4}$ | 0.035 | **0.289*** | $<10^{-4}$ | $<10^{-4}$ | $<10^{-4}$ |
| 2K | **0.571*** | 0.936* | 0.878* | $<10^{-4}$ | 0.001 | $<10^{-4}$ |
| 15K | **0.852*** | 0.898* | 0.888* | $<10^{-4}$ | 0.011 | $<10^{-4}$ |
| **Goodness-of-fit, Akaike's Information Criterion** | | | | | | |
| Background | Exponential | Gamma | Weibull | Normal | LogNorm | Logistic |
| Anc | -7032 | -7061 | **-7073** | -5802 | -6960 | -6134 |
| 2K | **-4873** | -4872 | -4376 | -4751 | -4872 | -4454 |
| 15K | **-3518** | -3516 | -3186 | -3449 | -3516 | -3233 |

[1]Note: Asterisks indicate satisfactory fit of distribution to the observed tail of beneficial mutations based on KS test ($p > 0.05$). Bold font indicates the best-fitting distribution based on AIC criterion for each genetic background.

**Table S3. Multiple linear model for prevalence of mutations in gene targets in Ara+2 population.**

| **Variable** | **Estimate** | **Std. Error** | **t value** | ***p*(>\|t\|)[1]** |
|---|---|---|---|---|
| Anc fitness | 73.130 | 23.510 | 3.111 | **0.002*** |
| 2K fitness | 82.300 | 25.070 | 3.283 | **0.001*** |
| 15K fitness | -19.490 | 20.050 | -0.972 | 0.332 |
| Target size | 0.003 | $< 10^{-3}$ | 12.176 | **$<10^{-15}$*** |

[1]Note: Asterisks and bold font indicate significant effects.

**Table S4. Primers used in this study.**

| Name | sequence |
|---|---|
| MS-custom-adaptor-F | 5'-TTCCCTACACGACGCTCTTCCGATCTNN-3' |
| MS-custom-adaptor-R | 5'-AGATCGGAAGAGCGTCGTGTAGGGAA-3' |
| HS-custom-adaptor-F | 5'-TCGTCGGCAGCGTCAGATGTGTATAAGAGACAGNN-3' |
| HS-custom-adaptor-R | 5'-CTGTCTCTTATACACATCTGACGCTGCCGACGA-3' |
| MS-junction-PCR-F | 5'-TCGTCGGCAGCGTCAGATGTGTATAAGAGACAGNNNNNNTTCCCTACACGACGCTCTTCCGATCT-3 |
| MS-junction-PCR-R | 5'-GTCTCGTGGGCTCGGAGATGTGTATAAGAGACAGNNNNNNAGACCGGGGACTTATCATCCAACCTGT-3 |
| HS-junction-PCR-F | 5'-TCGTCGGCAGCGTCAGATGTGTATAAGAGACAG-3' |
| HS-junction-PCR-R | 5'-GTCTCGTGGGCTCGGAGATGTGTATAAGAGACAGNNNNNNAGACCGGGGACTTATCATCCAACCTGT-3' |
